# Comparative transcriptomics of tropical woody plants supports fast and furious strategy along the leaf economics spectrum in lianas

**DOI:** 10.1101/2021.07.06.451334

**Authors:** U. Uzay Sezen, Samantha J. Worthy, Maria N. Umaña, Stuart J. Davies, Sean M. McMahon, Nathan G. Swenson

**Affiliations:** Smithsonian Environmental Research Center, 647 Contees Wharf Rd, Edgewater, Maryland, 21037 USA; Department of Evolution and Ecology, University of California, Davis, California, 95616 USA; Department of Ecology and Evolutionary Biology, University of Michigan, Ann Arbor, Michigan, 48109 USA; Forest Global Earth Observatory, Smithsonian Tropical Research Institute, Gamboa, Panama; Department of Botany, National Museum of Natural History, Smithsonian Institution, Washington DC, 20560 USA; Department of Biological Sciences, University of Notre Dame, Notre Dame, Indiana, 46556 USA

## Abstract

Lianas, climbing woody plants, influence the structure and function of tropical forests. Climbing traits have evolved multiple times, including ancestral groups such as gymnosperms and pteridophytes, but the genetic basis of the liana strategy is largely unknown. Here, we use a comparative transcriptomic approach for 47 tropical plant species, including ten lianas of diverse taxonomic origins, to identify genes that are consistently expressed or downregulated only in lianas. Our comparative analysis of full-length transcripts enabled the identification of a core interactomic network common to lianas. Sets of transcripts identified from our analysis reveal features related to functional traits pertinent to leaf economics spectrum in lianas, include upregulation of genes controlling epidermal cuticular properties, cell wall remodeling, carbon concentrating mechanism, cell cycle progression, DNA repair and a large suit of downregulated transcription factors and enzymes involved in ABA-mediated stress response as well as lignin and suberin synthesis. All together, these genes are known to be significant in shaping plant morphologies through responses such as gravitropism, phyllotaxy and shade avoidance.

## Introduction

Lianas are woody vines that have evolved multiple times since at least the Devonian period in vascular plant lineages, including in the Gnetales and repeatedly in the Angiosperms with fascinating diversity in stem anatomy even within a single genus [1,2]. The Neotropics carry close to 11,000 liana species belonging to 977 genera under 119 families [3], while the Old-world tropics contain about 12,000 species found in 143 families and 1415 genera [4]. In tropical forests, lianas may account for as much as 25% of the woody plant species, but can also represent 40% of canopy leaf cover [5, 6]. As structural parasites, lianas affect host tree recruitment, growth, survival, and reproduction [7–9]. Lianas also play significant roles in the cycling of carbon, nitrogen and water, and can hold back recovery after gap-forming events [6, 10–17]. Despite their exceptional diversity and ecological importance, little is known about the genetic characteristics that distinguish them from other vascular plants that help explain their convergence.

The basic advantage of the liana life form is clear: lianas harvest light from the treetops without investing in the woody structures necessary to support a canopy of leaves [18]. Several trade-offs emerge from this strategy of structural parasitism that might provide a path toward understanding convergent liana phenotypes. Specifically, in order to gain access to the canopy, lianas need to balance leaf area, length of vascular tissues, and tolerance to high temperature, excess light and dehydration which suggest potential underlying genetic mechanisms towards a liana growth form in multiple plant families [19, 20]. In the following, we consider each of these challenges to the liana lifestyle as a means of generating expectations regarding where similarities in gene expression profiles among a phylogenetically diverse set of lianas may exist. The liana life form involves a relatively longer stem length compared to that of trees culminating with a large total leaf area that sprawls over the crowns of many trees would make lianas more vulnerable to xylem vessel embolism than trees and shrubs [21, 22]. Liana leaves must have evolved adaptations that complement and compensate for hydraulic limitations and occupy a distinct range along the leaf economic spectrum (LES) [23, 24]. Compared to trees, lianas tend to have thinner blades with higher specific leaf area (SLA) and nutrient content. The foliar chemistry of lianas and trees also form a contrast based on leaf mass and area [25]. However, their leaves can also exhibit higher photosynthetic efficiencies despite the fact that the general leaf qualities reflect inexpensive, disposable, short longevity structures with little or no investment on defense [26].

The direct solar radiation and high heat in the canopies, where most liana leaves are deployed can impose significant constraints for lianas. For instance, the photosystem II has been shown to be vulnerable to temperature and light stress in a liana as compared to its tree relative [27]. Canopy trees have the potential to mediate this damage through higher leaf area with thinner boundary layers for efficient cooling. There is evidence that liana leaves have lower leaf transmittance values, better for light-capture, but worse for damage to tissues [27, 28]. It is highly possible that lianas have evolved certain epicuticular and parenchymal properties that have a set upper limit for safe light harvest and deflect excessive photosynthetically active radiation. When leaf temperature exceeds photosynthetic optimum, CO_2_ assimilation rates and stomatal conductance decrease while respiration increases [29]. Seasonal observations indicate that carbon fixation and efficiencies of water and nitrogen use are especially high in lianas during drought giving them a growth advantage [30–32]. We, therefore, expect that leaf surface qualities and mesophyll properties should be under selective pressures for dealing with light, thermal and dehydration stress. Guard cell openings can be set to maintain a steady rate of water loss enough to last through the day and from *Vitis vinifera* (grapevine from here in) we know that lianas can employ a range of anisohydric and isohydric control of stomatal conductance as water use strategy. Through their cheap but efficient leaves, lianas may also be carrying out sufficient levels of photosynthesis rapidly before midday to escape from the sun’s pernicious rays.

There has been some research on stress responses of a few liana species [33, 34]. However, most of the information comes from grapevine due to the large amount of genomic and transcriptomic information generated for the species [35–39]. A lack of high throughput sequencing data for additional liana species has reduced our ability to explore their potential genetic commonalities that distinguish them from non-liana plants.

Here, we provide a comparative transcriptomic analysis across a phylogenetically diverse group of trees, shrubs, and lianas. We use reference transcriptome assemblies of 37 tree and shrub species and ten liana species (SI Figure S1, SI Dataset S1). Using network analyses, we ask the following questions: (1) what genes are uniquely expressed and downregulated in some or all of the lianas? (2) Are there unique genetic similarities among lianas related to their shared ecological challenges (e.g., long distance nutrient transport, stress including hydraulic, thermal and light, leaf tissue quality, unidirectional growth and light sensing)? And (3) are there unique genetic similarities in lianas that reflect specific metabolic pathways underpinning the building of structural biomass?

## Materials and Methods

### Sample set

The Luquillo transcriptomic set included six lianas (DIO) *Dioscorea polygonoides* (Dioscoreaceae), (HET) *Heteropterys laurifolia* (Malpighiaceae), (PAU) *Paullinia pinnata* (Sapindaceae), (SEC) *Securidaca virgata* (Polygonaceae), (SMI) *Smilax coriaceae* (Smilaceae), (DOL) *Dolichandra unguis-cati* (Bignoniaceae). We also included four more liana species from National Center for Biotechnology Information’s Sequence Read Archive (NCBI-SRA) and One Thousand Plant Genomes (1KP) dataset [40, 41] with the following accessions (GNE) *Gnetum montanum* (Gnetaceae) SRR5908685, (SMISIE) *Smilax sieboldii* (Smilacae) SRR5134200, (PASCAE) *Passiflora caerulea* (Passifloraceae) 1KP id:SIZE, (SCHPAR) *Schlegelia parasitica* (Schlegeliaceae) 1KP id:GAKQ. A full list including non-liana species can be found in the Supplementary Information (SI Dataset S1, SI Figure S1). NCBI-SRA bioproject accession for the Luquillo transcriptomic set is PRJNA837288.

### Sample collection and RNA library construction

To analyze the transcriptomes, we chose healthy and fully developed leaves from seedlings of tree and liana species distributed between 350 and 450 m in elevation from the Luquillo Experimental Forest (LEF) in the north eastern part of Puerto Rico. For each species, approximately 5 grams of leaf tissue was collected and placed in a 50 mL polypropylene conical tube with RNAlater (Thermo Fisher Scientific, Waltham, MA, USA). Explants were cut with a razor blade prior to being placed in the tube to allow the RNAlater to penetrate the mesophyll quickly. Samples were then frozen at -80ºC within two days. Rneasy Plant Mini Kit (Qiagen, Valencia, CA, USA) was used for RNA extraction. RNA quantification and quality metrics were carried out using a NanoDrop 2000 spectrophotometer (NanoDrop Products, Wilmington, DE, USA) and an Agilent Bioanalyzer 2100 (Agilent Technologies, Santa Clara, CA, USA) RNAseq library preparations and sequencing were performed at the Beijing Genomics Institute, Shenzen, China on Illumina Hiseq 2000 sequencer generating 100 bp paired-end reads.

### Bioinformatics

Paired-end raw Illumina reads were trimmed, and quality filtered using Sickle (-q 35 -l 30 for minimum quality score and retaining sequences longer than 30 nt). Trimmed fastq files were assembled by Trinity v.2.6.6 with minimum contig length 300 [42]. Quality of the assemblies were assessed by QUAST (Quality Assessment Tool for Genome Assemblies), validated by Bowtie2, and analyzed by BUSCO for completeness using the Embryophyta (embryophyta odb9) benchmark gene set [43–45] (SI Dataset S1). Transcriptome assemblies were annotated by EnTAP using the Diamond high performance aligner interrogating four protein databases (Uniprot, RefSeq plant proteins 94, RefSeq complete protein 94, RefSeq non-redundant protein 94) [46, 47]. Bacterial, archaeal and non-plant eukaryotic contaminants were filtered using Opisthokonta as the taxonomical cut-off (SI Dataset S3). Transcriptomes were frame selected and translated into proteins through built-in GeneMarkS-T module within EnTAP [48] (SI Dataset S3). Full-length protein sequences were filtered from EnTAP results and were blasted against indexed *V. vinifera* proteome v29720. Top hits were selected through vsearch (-ublast -evalue 1e-9 -query cov 0.9) [49]. Blast results were matched into Grapevine UNIPROT IDs and used as input into STRING database for interactome network construction [50, 51]. Co-downregulated and co-expressed protein-protein interactome networks were imported into Cytoscape and merged into a single network [52] (SI Dataset S2). The resulting interactome network was annotated with colored borders and fills to highlight biologically informative nodes. Phylogenetic tree of lianas and non-lianas was constructed using the TimeTree resource compiling evolutionary divergence times derived from molecular sequence data [53]. Gene Ontology (GO) terms were interrogated using g:Profiler by providing gene lists corresponding to UNIPROT IDs [54]. Results generated from g:Profiler are accessible through permalinks in SI Text.

### Trait data

Leaf Area (LA) and Specific LA (SLA) trait data came from Zambrano et al. 2019 for mature trees in LEF [55]. For *Dolichandra unguis-cati* we used values from Osunkoya et al. (2014) [56]. Trait data for *Gnetum montanum* was obtained from the China Trait Database Wang et al. 2018 [57]. For the Luquillo vines *Dioscorea polygonoides, Heteropterys laurifolia, Paullinia pinnata, Securidaca virgata*, and *Smilax coriacea* we used unpublished data obtained from seedlings from our co-authors Samantha J. Worthy and Maria N. Umaña. Ordination of leaf traits was done by the ClustVis webserver [58]. Trait values and calculated PCA scores can be found in SI Dataset S1.

### A note on co-downregulation

Co-downregulation should not be confused with gene deletions observed in the genomes of many parasitic and carnivorous plants [59]. Absence of gene activity does not necessarily mean gene loss or loss of function. Therefore, here we prefer to use the term co-downregulation to define orthologous genes that are most likely present in lianas but not actively transcribed. Similarly, we define co-expression as orthologous genes from climbers demonstrating a sufficiently high expression pattern compared to non-liana counterparts that allows detection as full-length transcripts.

## Results

### Transcriptome assembly, annotation and trait comparison

Using a phylogenetically diverse sample of ten liana species, we identified major convergent gene expression patterns that contrast with 37 coexisting species of trees and shrubs (SI Figure S1, SI Figure S3). Quality metrics of transcriptomes generated by Quality Assessment Tools for Genome Assemblies (QUAST) showed the transcriptomes range in size from 49,040 (*Tetragastris balsamifera*) to 182,060 (*Ixora ferrea*) (SI Dataset S1). The overall alignment rates for the *de novo* assembled reference transcriptomes, validated by Bowtie2 alignments of trimmed and quality filtered reads, were between 83 and 94% (SI Figure S2A). The assembled transcriptomes captured an informative fraction of the expressed genes from 47 species where Benchmarking Universal Single-Copy Orthologs (BUSCO) analyses were able to represent between 52 and 92% of the benchmark eukaryotic genes within the Embryophyta (SI Figure S2B, SI Dataset S1). Annotation and filtering of non-plant transcripts through EnTAP (Eukaryotic non-model Transcriptome Annotation Pipeline) resulted in a full-length set of grapevine orthologs ranging from 6001 (*Smilax coriacea*) to 9488 (*Dolichandra unguis-cati*) for cross-comparison (Table 1, SI Dataset S3). An ordination of key LES traits, leaf area (LA) and specific leaf area (SLA), reflected difference between lianas and non-lianas in our transcriptome set, indicating a phenotypic portrayal of liana biology at a local island scale (SI Figure S4).

**Table 1.**
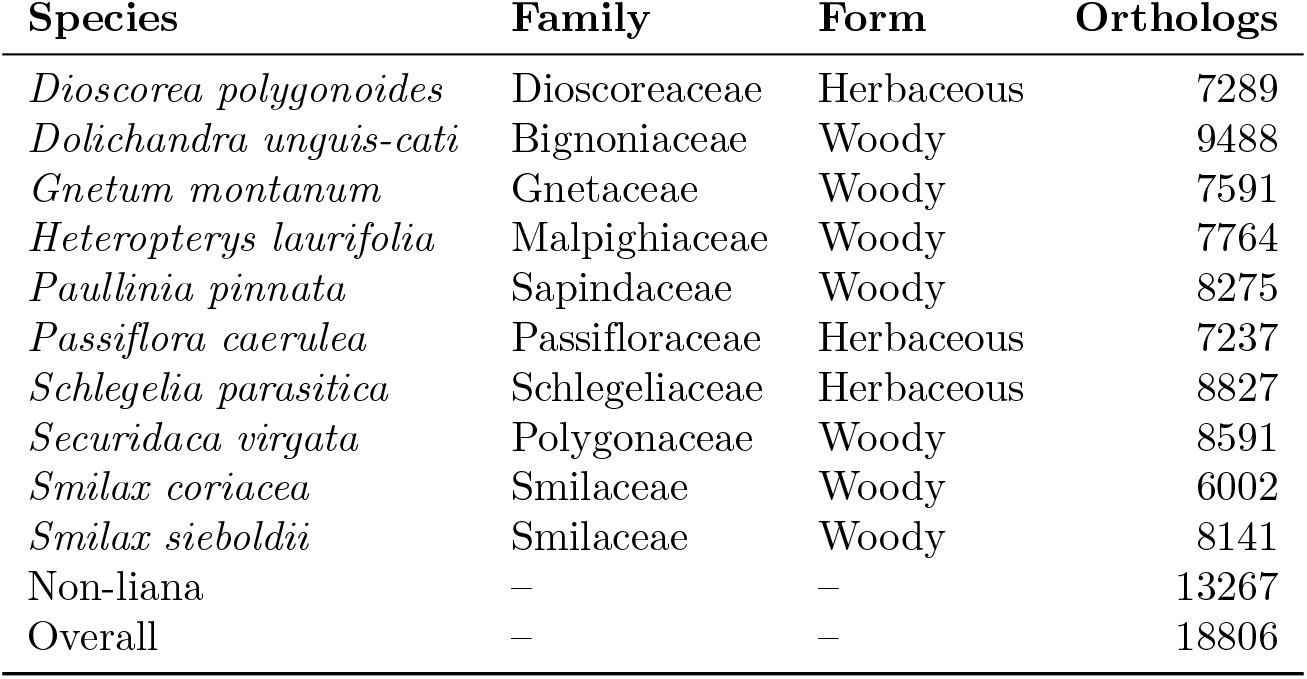
Ortholog numbers, families, and growth forms of species with full-length transcripts BLASTed onto grapevine protein models.

### A protein-protein interactome overlaid upon transcriptomic base layer

A Venn diagram of grapevine protein orthologs corresponding to our transcripts revealed 827 protein models common to all 47 plant species (SI Dataset S1). After the exclusion of common proteins, a protein-protein interactome network based on curated STRING database was constructed using transcripts exclusively expressed by lianas and non-lianas. The total size of the network including vertices with first and second shells of interaction consisted of 3835 nodes and 14574 edges. Grapevine has 33,568 protein-coding genes and the network occupies a little over 11% of its gene space. Close to half of the network (1687 nodes) comprised genes solely expressed by non-lianas (Fig. 1). The rest included transcripts exclusively expressed by one or more lianas (Fig. 1A). The most connected node (VIT 11s0016g03430) with 199 interactions represented the core of the interactome corresponded to a protein phosphatase 2C (PP2C) also known as cyclic nucleotide-binding/kinase domain-containing protein (SI Figure S3). This protein-serine/threonine phosphatase is a key enzyme in cGMP-dependent signaling, abscisic acid perception, commitment to cell cycle and is found within the additional shells of interaction in a region of the network subtending nodes exclusively expressed in lianas (Fig. 1, Table 2, SI Figure S3). The subsequent highly connected nodes are all found within the co-downregulated portion of the network (Fig. 1B, Table 2)

**Figure 1.**
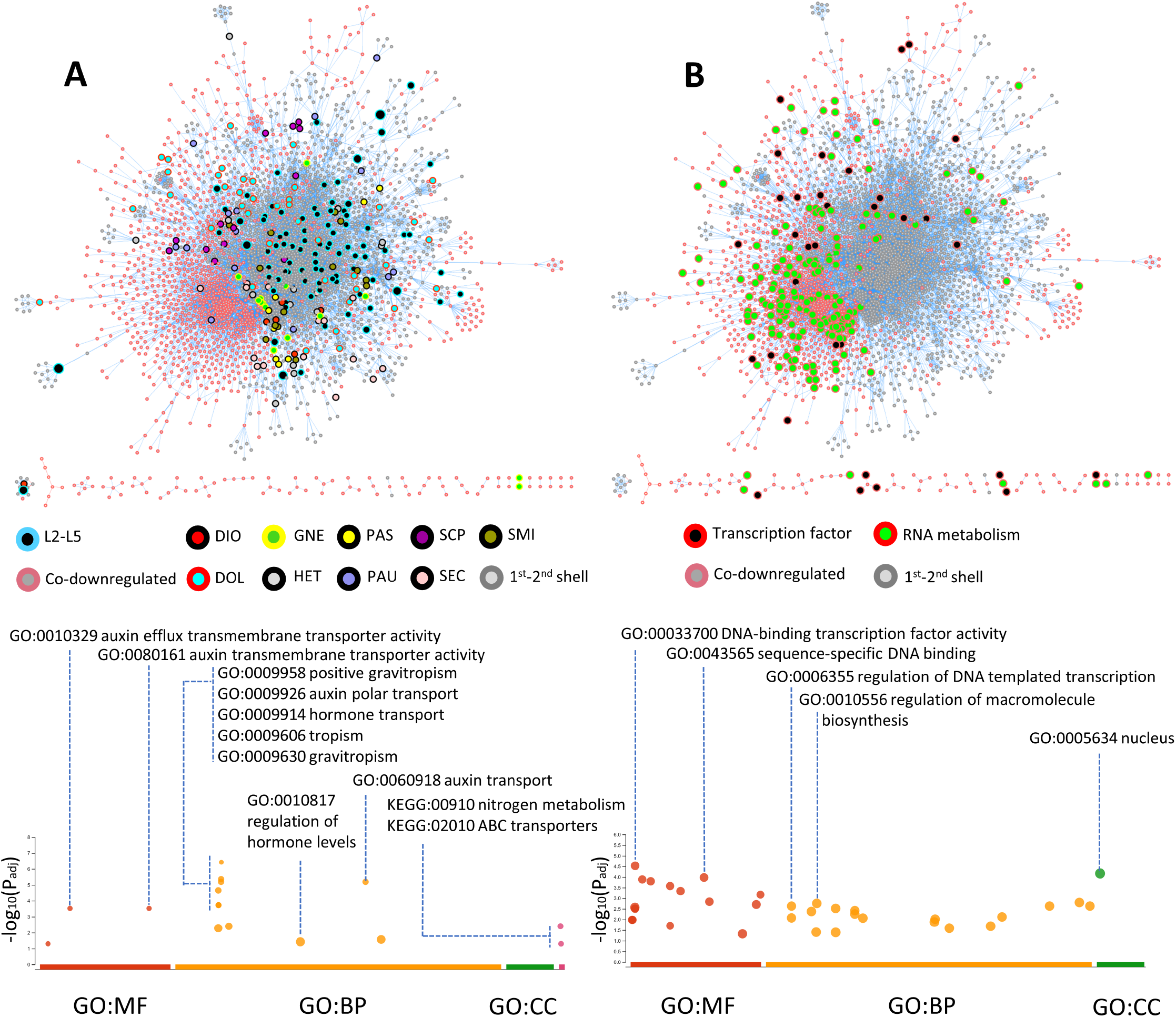
Protein orthologs of full-length liana and non-liana transcripts mapped onto grapevine interactome and Manhattan plots of enriched GO terms. **A**, Protein-protein interactions of transcripts solely expressed in ten lianas (the two Smilax species are represented as combined into SMI). Nodes shared by more than two lianas (L2-L5) and co-downregulated transcripts (salmon-colored borders) including their first shell interactors (gray colored borders) occupy distinctly sectorized footprints. GO terms represented molecular functions (MF) pertaining to the phytohormone auxin, biological processes (BP) including positive gravitropism and auxin transport. KEGG pathways included nitrogen metabolism and ABC transporters. No enrichment observed in cellular compartment (CC). **B** Nodes pertaining to mRNA metabolism (red border and green fill) and transcription factors (red border and black fill) are observed to be downregulated in all lianas in the dataset. GO term analysis of co-downregulated transcripts revealed terms such as regulation of gene expression concentrated around transcription factors. Molecular functions were enriched in DNA-binding transcription factor activity and sequence-specific DNA binding. Enriched biological processes included regulation of DNA templated transcription and regulation of macromolecule biosynthesis all localized to the nucleus.

**Table 2.**
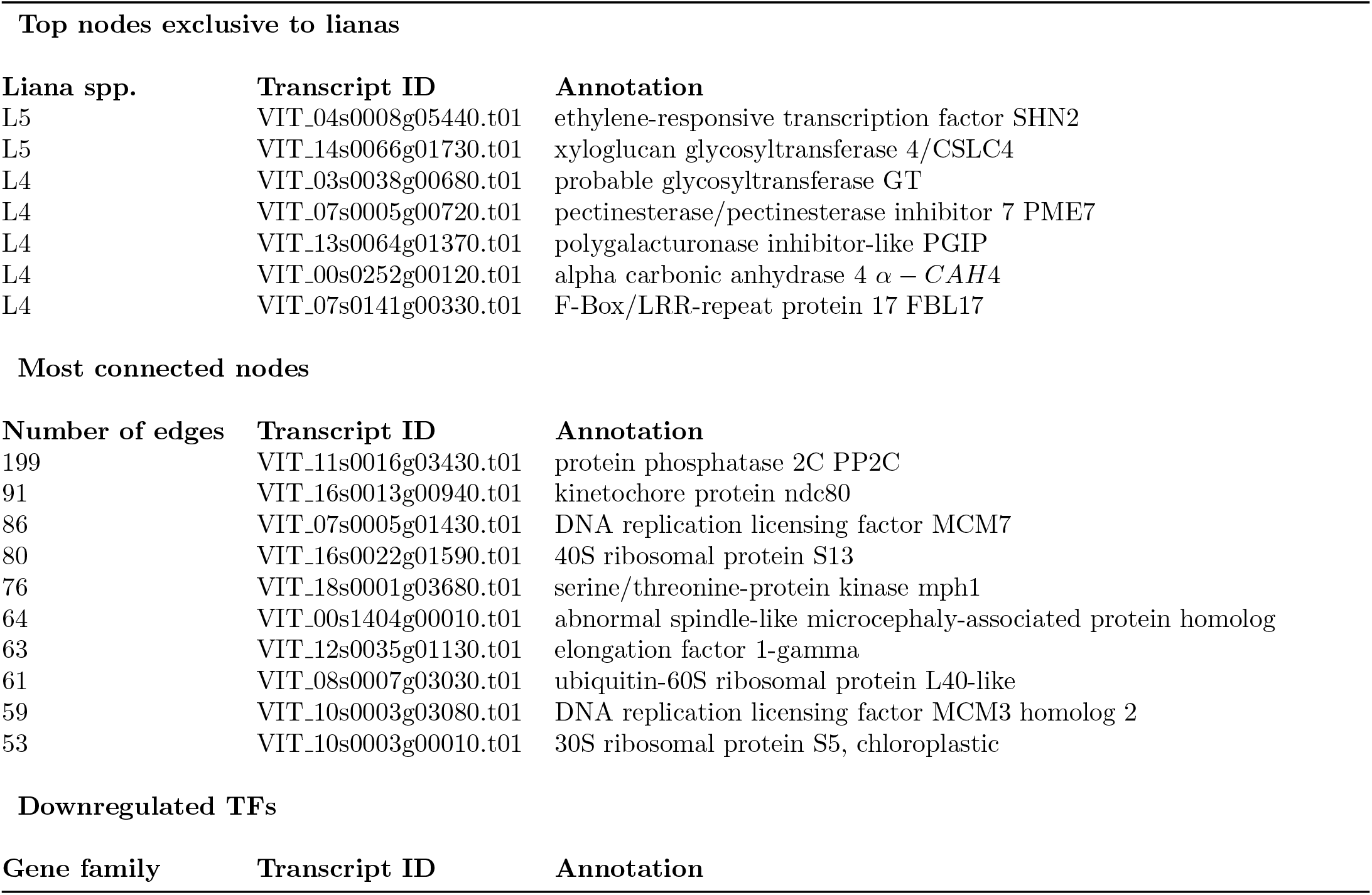
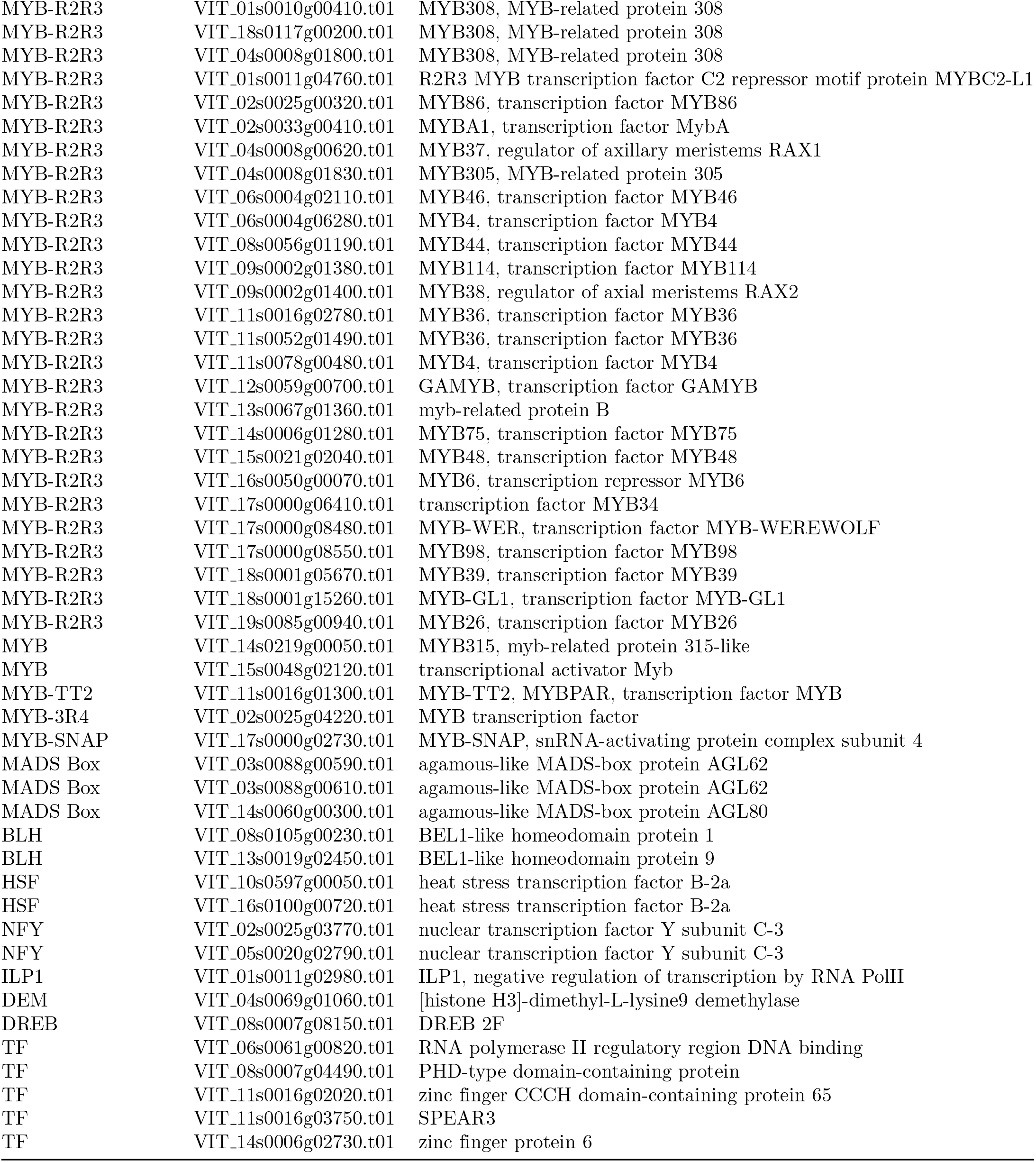
Description of nodes of interest including transcripts exclusively expressed by more than one liana species (L2-L5), most connected nodes, and transcription factors (TFs) found within the co-downregulated portion of the protein-protein interactome network.

### Transcripts exclusively expressed in lianas

The transcripts expressed exclusively by more than one liana (L2-L5) species generated a network of 705 nodes with 1609 edges (Fig. 1, SI Figure S3). We found no genes that are unanimously expressed by all lianas. Two nodes, shared by five lianas (L5), corresponded to the ethylene responsive transcription factor SHINE2 (SHN2) and the xyloglucan glycosyltransferase 4/cellulose synthase-like C family 4 (CSLC4) enzyme, representing the highest points of convergence via individual gene transcripts (Table 2). The next set of most nodes shared by four lianas (L4) in different combinations included cell wall remodeling enzymes pectin methylesterase 7 (PME7), glycosyl transferase (GT), polygalacturonase inhibitor-like (PGIP) as well as a carbon concentrating enzyme alpha carbonic anhydrase 4 (alpha-CAH4), and the plant-specific cell cycle regulator F-box-like 17 (FBL17) (Table 2, Figure S3).

The combined controlled vocabulary gene ontology (GO) analysis of transcripts shared by more than two lianas (L2-L5) through gProfiler displayed enriched molecular functions pertaining to the phytohormone auxin, such as auxin efflux transmembrane transporter activity (GO:0010329) and auxin transmembrane transporter activity (GO:0080161) (Fig. 1A). Biological processes included auxin associated activities such as positive gravitropism (GO:0009958), auxin polar transport (GO:0009926), hormone transport (GO:0009914), auxin transport (GO:0060918), tropism (GO:0009606), response to gravity (GO:0009629), gravitropism (GO:0009630) and regulation of hormone levels (GO:0010817) (Fig. 1A). Enriched Kyoto Encyclopedia of Genes and Genomes (KEGG) pathways included nitrogen metabolism (KEGG:00910) and ABC transporters (KEGG:02010) (Fig. 1A).

The term auxin and gravitropic response identified by gProfiler were consolidated around five genes including chaperone protein dnaJ 15-like (VIT 01S0011G03790), ABC transporter B family member 1 (VIT 08S0007G05060), protein kinase PINOID-like (VIT 10S0003G04320), auxin efflux carrier component (VIT 11S0052G00440), SEC7 domain-containing protein (VIT 02S0012G01790), and auxin efflux carrier component (VIT 14S0108G00020) (SI Dataset S1).

### Transcripts not expressed in lianas

Comparative analysis revealed 2684 grapevine orthologs that were not expressed in our liana transcriptome set (SI Dataset S1). Of these, 1687 had hits in the STRING interactome database (Fig. 1). The co-downregulated transcripts (i.e. transcripts that probably have a corresponding gene copy in the liana genomes but were not detected in our set of liana transcripts) enveloped a large portion of the network with 3751 edges. GO term analysis through gProfiler revealed terms pertaining to regulation of gene expression highly concentrated around transcription factors. Molecular functions were enriched in DNA-binding transcription factor activity (GO:0003700) and sequence-specific DNA binding (GO:0043565). Enriched biological processes included regulation of DNA templated transcription (GO:0006355) and regulation of macromolecule biosynthesis (GO:0010556), which are localized to the nucleus (GO:0005634) (Fig. 1B). Transcription factors such as nuclear factor Y (NFY), agamous-like (AGL), dehydration responsive element binding (DREB), BEL-like homeodomain (BLH), and myeloblastosis (MYB) showed significant presence in the co-downregulated sector of the interactome. The largest group of transcription factors consisted of MYB family represented by 32 members where 27 of them belonged to MYB-R2R3 sub-family (Table 2). The co-downregulated portion of the interactome is comprised of nodes pertaining to mRNA metabolism (Fig. 1B).

### Comparison of three families having liana and non-liana pairs

Three families belonging to Bignoniaceae, Sapindaceae, and Malpighiaceae included pairs of liana and non-liana members in our dataset. We scrutinized these with the anticipation of detecting the most informative gene expression differences (Fig. 2). Enrichment analysis for transcripts expressed only in the liana members resulted in five GO terms and a single KEGG pathway shared between Bignoniaceae and Sapindaceae. These shared terms are involved in developmental process during reproduction (GO:0003006), system development (GO:0048731), intracellular anatomical structure (GO:0005622), chloroplast (GO:0009507), plastid (GO:0009536), and biosynthesis of cofactors (KEGG:01240) (Fig. 2). Terms enriched in Malpighiaceae did not show any overlap with the other two families (SI Dataset S1).

**Figure 2.**
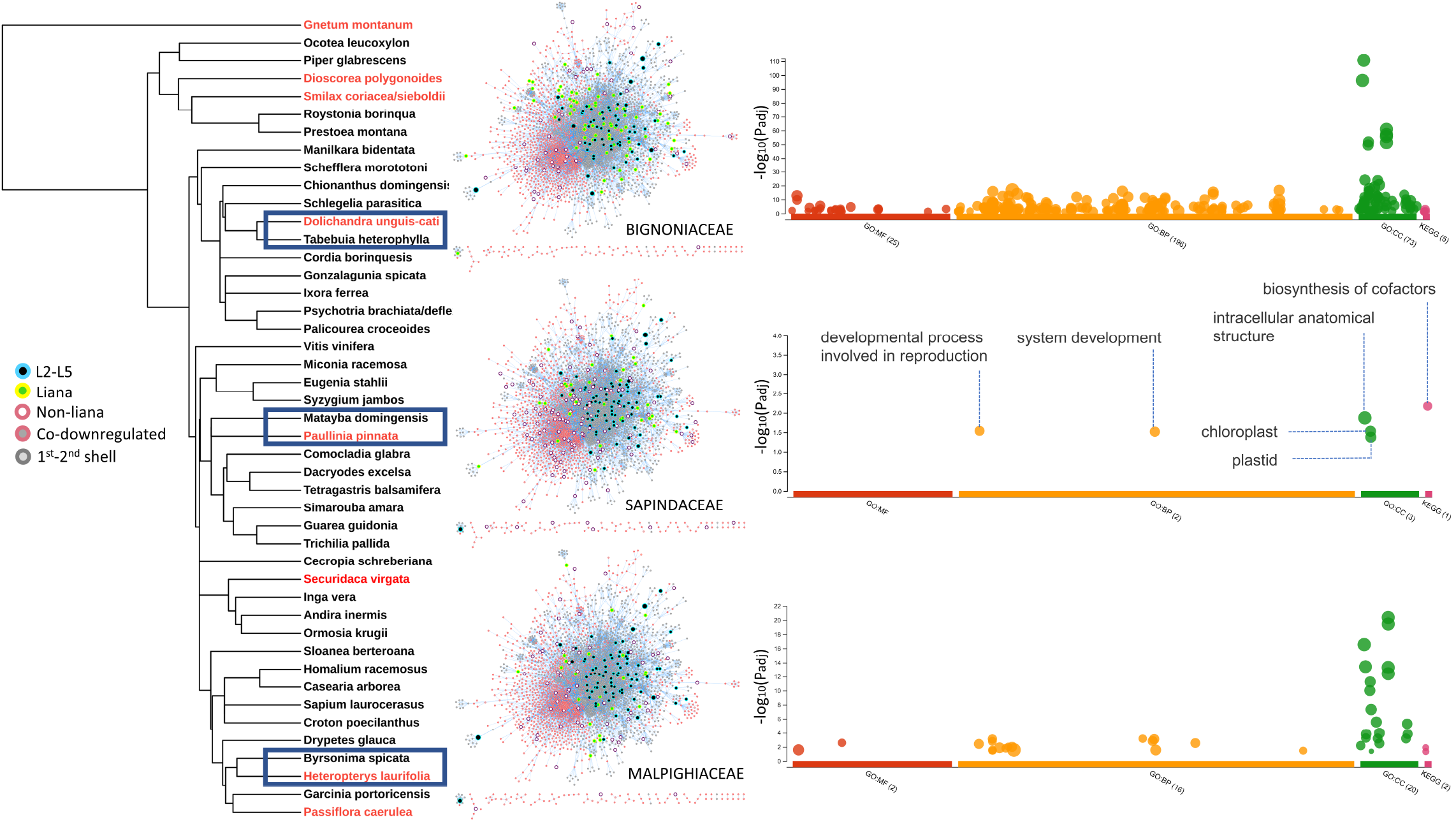
Phylogenetic placement of lianas and non-lianas together with interactome and GO term enrichment of three families bearing liana and non-liana pairs: *Dolichandra unguis-cati* vs *Tabebuia heterophylla* (Bignoniaceae), *Paullinia pinnata* vs *Matayba domingensis* (Sapindaceae), *Heteropterys laurifolia* vs *Byrsonima spicata* (Malpighiaceae). Members of the lianas (yellow colored border with green fill) and non-lianas (salmon colored border with no fill) are highlighted inside interactome networks for each of three within-family comparisons. Co-expressed nodes shared by more than two lianas (L2-L5) are retained for visual guidance as in Fig. 1. A total of six enriched GO and KEGG terms were shared between *D. unguis-cati* and *P. pinnata*. Terms enriched in *H. laurifolia* did not show overlap with the other two lianas. Permalinks for the gProfiler GO term analysis are *P. pinnata* vs *M. domingensis*: https://biit.cs.ut.ee/gplink/l/TZwIGOGgSk; *H. laurifolia* vs *B. spicata*:https://biit.cs.ut.ee/gplink/l/FmLAWHzES9; *D. unguis-cati* vs *T. heterophylla*:https://biit.cs.ut.ee/gplink/l/iXBz-L_6_*T*2

## Discussion

Lianas and trees represent two widely separate growth forms with distinct trade off in biomass allocation and growth. Instead of self-supporting stems, lianas allocate most of their resources on shoot and leaf production. Liana leaves must have evolved adaptations to reduce hydraulic failure due to vascular embolism, damage on light harvesting complexes due to intense light and thermal stress, dehydration due to insufficient control of stomatal conductance, inefficient carbon fixation due to mesophyllic carbon diffusion limitations. Here, we provide a wide snapshot of physiologically interpretable differences in gene expression between lianas and non-lianas within a subtropical plant community and make an attempt to learn about genotypic drivers of life-history phenotypes from transcriptional responses of co-habitants. [60–62] (SI Figure S1, SI Dataset S1). At early developmental stages, transcriptomic differences between woody lianas and herbaceous vines are most likely minimal. For this reason, we included three herbaceous climbers in the dataset with the assumption that leaves of woody and herbaceous climbers would experience similar constraints independent from their stem morphology and would perform in parallel (Table 1). Despite the ontogenetic and tissue diversity limitations, there exists a suite of differences consistent with expectations from the biology of climbing plants and functional traits theorized in LES [23, 24, 26, 63, 64]. Top leaf level functional traits of lianas correspond to higher SLA and maximal photosynthetic rates. Enzymes we highlight orchestrate cuticular and parenchymal properties related to mesophyll conductance (Fig. 3). Liana leaves may do more with less through selective endoreduplication. The resultant action of cell wall building enzymes and control of lignification appears to be in line with the predictions of the LES in lianas. Downregulation of a high number of MYB transcription factors belonging to the largest gene family in plants indicates a crosstalk with cell wall building enzymes since many R2R3-MYBs are known to be involved in lignin deposition. A few R2R3-MYBs are also involved in ABA-mediated control of transpiration which may explain growth advantage experienced by lianas observed in seasonally dry tropical forests [31, 32, 65].

**Figure 3.**
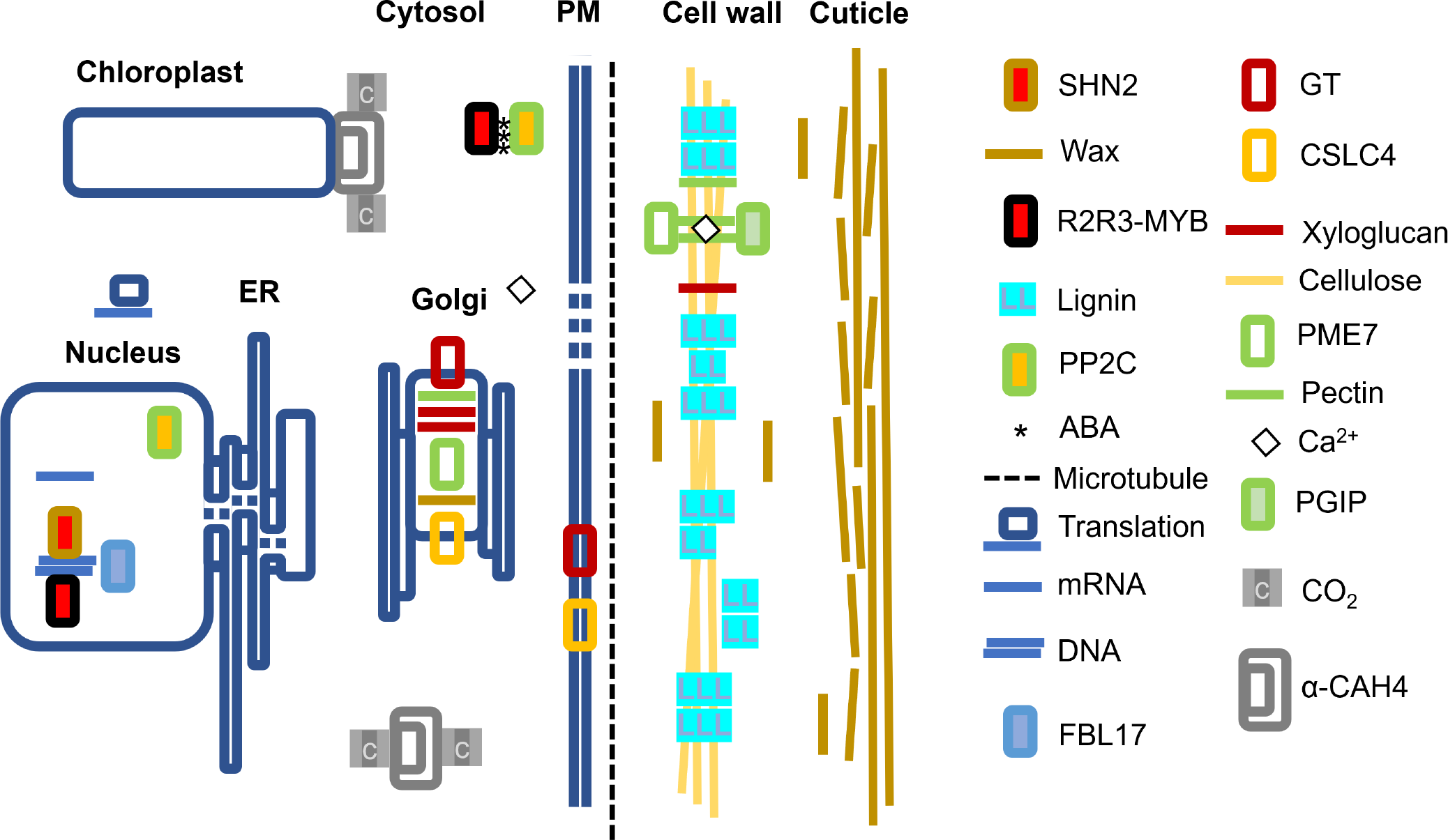
A conceptual sketch of transcripts and their cellular compartments highlighted in this study. Lianas appear to invest in thickened cuticular layers through upregulation of SHN2. Cell wall remodeling enzymes CSLC4, GT, PGIP, PME7 similarly show upregulation where CSLC4, PME7 and GT are Golgi localized together with pectin, xyloglucan and wax. GT and CSLC4 move into plasma membrane (PM) and deposit xyloglucan, pectin, and cellulose onto the cell wall following microtubule orientation patterns. Upregulated FBL17 is involved in cell cycle control, endoreduplication and double-stranded DNA damage response. Alpha-CAH4 is localized to the cytosol and chloroplast and is a significant player in the CO_2_ concentrating mechanism. A large number of R2R3-MYB transcription factors were co-downregulated in lianas playing significant roles as activators and repressors controlling diverse functions such as pectin, lignin and suberin mediated xylogenesis, branching and phyllotaxis and ABA mediated stress tolerance. Cell wall qualities are affected by lignin deposition and pectin crosslinking complexed with Ca^2+^. PP2C is the most connected node forming the core of the interactome as a key player in ABA induced stress response and interacts with MYBs. Salient liana leaf transcripts collectively may form genetic features underlying the fast end of the LES.

### Top co-expressed transcripts in lianas include enzymes relevant to LES

Top co-expressed genes in our analysis correspond to enzymes synthesizing or modifying cell wall components, cuticle properties, carbon concentrating mechanism, cell cycle progression through endoreduplication and double-stranded DNA damage response (Table 2, Fig. 3, SI Figure S3). Modifications of cell wall components and cuticle properties are beneficial for lianas in developing leaves with high SLA and optical properties since excessive light levels may damage light harvesting complexes. Reduced production cost due to altered secondary cell wall composition, increased rates of carbon fixation and fast tissue turnover in most liana species may be an outcome of series of mesophyllic adaptations. Facilitation of carbon capture and diffusion through an active carbon concentrating enzyme capable of scavenging low levels of CO_2_ floating inside the leaf could be adaptive since leaves with high SLA can accommodate only a few mesophyll cell layers. Lianas could be achieving more with less by increasing gene dosage through selective endoreduplication. In lianas, under hydraulic constraints, machinery evolved for the repair of damaged nuclear genetic material may be primed as early as the seedling stage.

### Cuticle properties controlled by SHN2

We found co-expression of SHN2 transcription factor exclusively by five liana species involved in regulation of lipid biosynthesis (Table 2, Fig. 3, SI Figure S3). SHNs together with wax inducers (WINs) stimulate biosynthesis of cuticular wax in leaf epidermis and reduce stomatal density [66, 67]. SHNs can induce changes in the leaf cuticle to make it more resistant to light stress and avoid excessive heating [68]. Overexpressed SHN2 may indirectly suppress the biosynthesis of lignified secondary wall [69]. This is relevant for lianas because the leaf economical spectrum of lianas predict forms with cheap, disposable but efficient blades with reduced longevity and defense [26]. In our study, many MYBs, some of which involved in lignified secondary wall biosynthesis, have been observed to be downregulated in lianas (Fig. 1B, Table 2, SI Figure S3). All 149 genes belonging to AP2/ERF protein family in grapevine genome have been transcriptionally characterized and the Arabidopsis SHN2 ortholog gene (GSVIVP00032652001/VvERF044) also showed an upregulated pattern in the grapevine leaves [70].

### Cell wall remodeling enzymes

We observed upregulation of a suite of cell wall remodeling enzymes such as CSLC4, PME7, GT, PGIP (Fig. 1A, Fig. 3, Table 2, SI Figure S3) [71]. These cell wall modifying enzymes not only determine leaf morphogenesis but also phyllotaxis [72]. Less lignified leaves with altered cellulose and hemicellulose composition may be one way to achieve set of traits defining liana leaves with fast growth, resource uptake and high productivity end of the leaf economic spectrum. The rigidity of the cell wall is proportional to the number of xyloglucan connections maintained through Golgi-located xyloglucan synthesizing enzymes [73, 74]. Removal of xyloglucan ties loosens cell wall leading to asymmetries in elasticity and resulting in highly convoluted structures [75]. Overexpression of xyloglucanase in transgenic poplar resulted in droopy leaves with petioles slanted downward [76]. This could be significant because lianas tend to evolve longer petioles when exposed to full light and rigidity of petioles can determine photosynthetic efficiency especially in cordate leaves [77, 78]. Microtubules laid on the cell wall during development can become a reinforcing factor in anisotropic growth since cellulose synthases move along those tubules altering orientations of cellulose deposits [79]. Cellulose is the main load bearing component especially in secondary cell walls [71, 80]. Quintuple mutant analysis of Arabidopsis has revealed that besides CSLC4 four other CSLC genes (CSLC5, CSLC6, CSLC8, and CSLC12) also possess xyloglucan synthase activity [81]. Another cell wall remodeling enzyme showing upregulation among lianas is the PME7 (Table 2, Fig. 3, SI Figure S3). A composition of esterified and de-esterified pectin residues control calcium-mediated stiffness of gel matrix within the cell wall and contributes to diversification of organ shapes through directional growth. Arabidopsis mutants show that xyloglucans and pectin can together influence shoot meristem and phyllotaxis [72]. Cell wall remodeling could be a morphogenetic response resulting in shade avoidance in lianas [81]. For instance, disrupted phyllotaxy observed in Arabidopsis mutants may have been reinforced in lianas by coordinated expression of cell wall modifying enzymes which are also responsible from gravitropic response such as the positive gravitropism (GO:0009958), tropism (GO:0009606), response to gravity (GO:0009629), gravitropism (GO:0009630) enriched among GO term analysis of upregulated transcripts (Fig. 1A) [72, 73].

### An active component of carbon concentrating mechanism along the mesophyll may increase fixation efficiency of liana leaves

Our results also show exclusive activity of alpha-CAH4 in lianas but not in trees and shrubs indicating enhanced carbon concentrating mechanism along the liana mesophyll (Table 2, Fig. 3, SI Figure S3). The carbon concentrating mechanism describes the enzymatic pathways the atmospheric CO_2_ flows through the mesophyll cell wall and the plasma membrane into the liquid cytosolic phase imposing limitations on diffusion known as mesophyll conductance. Mesophyll conductance is a significant driving factor in LES [64]. Carbonic anhydrases are zinc metalloenzymes driving the reversible hydration of CO_2_ significantly aiding downstream carboxylases such as pyruvate carboxylase, acetyl-coenzyme A carboxylase, PEP carboxylase and RUBISCO [82]. Plants have three classes (alpha-, beta-, gamma-) of carbonic anhydrases [83]. Alpha-carbonic anhydrases are the largest class in the plant kingdom and are highly compartmentalized [84]. Their expression has been detected in cytosol and chloroplast. Carbonic anhydrases can capture CO_2_ generated by dark respiration as well as photorespiration enhancing carbon assimilation efficiency of RUBISCO. Stomatal metrics and morphology can determine water use efficiency [85]. Liana leaves with typically lower stomatal densities could be expressing alpha-CAH4 in higher levels increasing the efficiency of the spongy mesophyllic conductance along the CO_2_ diffusion pathway (Fig. 3). An active carbonic anhydrase may contribute to the maintenance of lower stomatal densities. Fossil evidence and present-day observations confirm that plants grown in elevated CO_2_ environments decrease their stomatal conductance and stomatal index [86, 87]. Plants over-expressing carbonic anhydrase in their guard cells have the potential to improve their water use efficiency. In guard cells, carbonic anhydrases serve as upstream regulators induced by CO_2_ levels independent of leaf photosynthetic rates [88]. High carbonic anhydrase activity could be making lianas more resilient in hotter and drier conditions by reducing the need for evapotranspiration [32, 65].

### Liana leaves express FBL17 transcription factor controlling cell cycle progression and double-stranded DNA damage response

We have observed that four lianas exclusively expressed the plant-specific FBL17 transcript that serves as a major checkpoint in cell cycle progression from Gap1 (G1) into Synthesis (S) phases [89] (Table 2, Fig. 3, SI Figure S3). From the LES perspective, leaf construction is the sum of spatially controlled cytoplasmic growth and cell division. During leaf expansion, a plant can do more with less through a process called endoreduplication where distinct cells may increase their genomic content and gene dosage without cell division [90–92]. During G1 into S phase transition cyclin dependent kinase type A (CDKA) and cyclin D (CYCD) work together to trigger an expression cascade of DNA replication genes. FBL17 is an indispensable protein component of E3 ubiquitin ligase SCF (ASK-Cullin-F-box) for ubiquitination of CDK inhibitor proteins KRPs (KIP-related protein/inhibitor-interactor of CDKs) for proteasomal degradation and forms a checkpoint in G1/S transition together with CDKA/CYD complex. Unregulated FBL17 expression causes disruptions in cell cycle genes, while at the same time it can be involved in double stranded DNA damage response leading to abnormalities in root and shoot meristems [93, 94]. In plants, FBL17 is involved in double stranded DNA break induced damage response, which is crucial for cell cycle arrest at the first G1/S checkpoint before being committed to division [94, 95]. Loss of function in FBL17 shuts off endoreduplication in Arabidopsis trichomes [93]. Liana leaves appear to be well protected especially against UV-induced damage with thickened secondary cell wall and waxy cuticle, therefore, upregulation of FBL17 might be a deliberate strategy to avoid interruptions in cell cycle. Downregulation of MYB4 and MYB75 involved in biosynthesis of UV-protecting flavonoids and anthocyanins and MYB46 involved in upstream of endoreduplication regulators could be a similarly reconciliatory exertion [96–99] (Table 2, SI Figure S3). Compatible with FBL17, some of the enriched GO terms for the top ten most connected nodes include (GO:0022402) cell cycle, (GO:0007049), double-strand break repair via break-induced replication (GO:0000727) (SI Dataset S1).

### Co-downregulated transcripts are dominated by MYB-R2R3 transcription factors

Silencing of even a single transcription factor can lead to drastic changes in phenotype with many examples from domesticated plants [100, 101]. In our comparisons, transcripts not expressed in lianas are overwhelmingly dominated by R2R3 subclass MYB transcription factors (Fig. 1, Fig. 3, Table 2, SI Figure S3). Significance of MYB-R2R3 co-downregulation for lianas could include stomatal control, lignification and plant architecture including phyllotaxis. The MYB-R2R3 serve as major activators and repressors controlling diverse functions such as pectin, lignin and suberin mediated xylogenesis through control of the phenolic acid metabolism, modulation of developmental signaling, cell cycle, epidermal cell fate and patterning [102–105]. There may be a crosstalk among the silent MYB-R2R3 transcription factors and exclusively upregulated set of cell wall building enzymes together shaping the liana leaf economy.

MYB genes are found in all eukaryotes and is the largest transcription factor family in Arabidopsis [105]. The MYB-R2R3 subfamily in land plants (Embryophyta) has expanded during the Silurian widening the structural complexity of tissues through secondary cell wall biosynthesis and vascularization [106–108]. Functional predictions of most MYBs have been sufficiently characterized both experimentally and computationally in myriad model plants. The grapevine and Arabidopsis have 134 and 126 R2R3-MYB genes, respectively [105, 109]. Structurally, the MYB family harbors significant levels of intrinsically disordered regions outside their canonical DNA-binding domains that possess potential for other functions regulated by post-transcriptional modifications [110]. Here we will highlight a subset of the R2R3-MYBs (MYB44, MYB4/MYB308, MYB46, MYB39, MYB38, MYB37) showing no expression in lianas (Fig. 1, Table 2, SI Figure S3).

### MYB44 and ABA-mediated stress tolerance

One particular R2R3-MYB downregulated in our analysis is MYB44 that deserves special attention since it is heavily involved with the core node of the interactome the protein phosphatase PP2C (Table 2, SI Figure S3). Lianas adopted features such as smaller and low-density stomatal openings with sunken guard cells for reduction of water loss due to excessive transpiration as early as the Paleozoic based on cuticular analysis of pteridosperm lianas [111]. The sesquiterpenoid plant hormone ABA mediates crucial physiological processes optimizing water use, carbon uptake and light signaling. Although guard cells can endogenously synthesize ABA, the main source for the whole plant is vascular parenchyma. In the presence of ABA, ABA receptors (PYR/PYL/RCAR) aggregate with clade A PP2Cs reducing the ability of PP2Cs to inhibit sugar non-fermenting 1 related protein kinase 2 (SnRk2) kinases. Freed from PP2C control, SnRK2 kinases phosphorylate and arm transcription factor genes to collectively induce or repress ABA-responsive genes downstream elucidating ABA-dependent stress response [112, 113]. The behavior of MYB44 appears to have opposing modes. In one mode, under stress MYB44 exerts suppressive effect on PP2C. The over expressed MYB44 has been shown to have antagonistic interaction with PP2C enzymes which control ABA signal transduction cascade in concert with ABA receptors [112, 114, 115] (Table 2). Overexpressed MYB44 also appeared to have binding interaction with ABA receptors [116]. In the other mode, MYB44 binds to transcription start sites of PP2C genes and represses their expression in normal unstressed conditions [117]. Moreover, akin to the prokaryotic end-product repression, MYB44 can bind to its own promoter in an act of self-silencing [118]. Induction of PP2Cs is reported from drought challenged grapevine transcriptomes [119]. In lianas, absence of MYB44 activity may be releasing repressed PP2C and could be a desensitization mechanism for less aggressive stomatal control. For instance, transgenic gray poplar trees expressing mutant ABA insensitive 1 (abi1) belonging to PP2Cs led to large and unresponsive stomata with inhibited lateral bud growth [120]. Downregulation of MYB37 could be a part of this desensitization scheme [121] (Table 2, SI Figure S3).

One noteworthy observation is that the top connected node forming the core of our interactome is another PP2C with kinase activity belonging to clade L protein phosphatases (VIT 11s0016g03430.t01/AT2G20050) [115](Table 2, SI Figure S3). The secondary messenger cyclic GMP binds and inhibits the phosphatase activity of this PP2C in favor of kinase activity. For this reason, this PP2C is known as cGMP-dependent protein kinase (PKG) and it phosphorylates the transcription factor GAMYB to upregulate gibberellic acid-responsive genes [122]. Leaf cell expansion following cell division also revolves around PP2C through ATPase activity acidifying cell walls [92]. In addition to PP2C, many of the rest of the highly connected nodes are pertaining to chromatin modification (Table 2). Chromatin modelers modify histone tails and contribute to the repressor activity of MYB44 ([117, 118].

MYB44 is central in transduction of multiple local abiotic stresses into whole plant response. For instance, SHN2 and MYB86 are responsive to excessive light stress in leaves but when plants experience heat stress simultaneously, incorporation of multiple forcings into a systemic acclimation is carried out by mediators including MYB44 [68].

### R2R3-MYBs as regulators of secondary cell wall biosynthesis

Lignin biosynthesis is repressed and activated by several R2R3-MYB genes, and it is curious whether the downregulated R2R3-MYBs identified in our study show biological associations with the set of upregulated cell wall remodeling and epicuticular enzymes (Fig. 1, Table 2, SI Figure S3). For instance, overexpressed SHN2 suppresses many MYBs involved in lignified secondary wall biosynthesis as evidenced from poplar [69]. Transcriptional regulation of secondary cell wall related lignin deposition in liana leaves appears to be fine-tuned since they downregulate a set of R2R3-MYBs serving as suppressors and activators of lignin biosynthesis [80] (Table 2). Lignin biosynthesis contains many redundant enzymes and lianas may be regulating multiple check points through silencing of multiple MYBs to achieve their functional leaf morphology encapsulated by LES hypothesis [64, 99].

### Downregulated R2R3-MYBs as repressors of lignin biosynthesis

We have identified MYB46, MYB4/MYB308, and MYB75 reported to be involved in lignin repression downregulated in liana leaves [123] (Table 2, SI Figure S3). MYB46 targets a set of 13 genes in lignin biosynthesis [80, 99, 123–125]. A repressor of endoreduplication called E2Fc is also known to bind to the promoter of MYB46 and suggests a potential crosstalk with the cell cycle regulator FBL-17 [99, 124]. Overexpressed MYB4, an ortholog of MYB308, leads to suppression of lignin biosynthesis and reduced elongation of stem internodes, which is a characteristic liana-associated trait [126]. MYB4 is induced by MYB46 but also (similar to MYB44) is inhibited by its own in an end-product repression fashion [80, 123]. MYB75 is a repressor of lignin and secondary wall biosynthetic genes but also an inducer of anthocyanins and flavonoids. Loss of function mutants in Arabidopsis displayed thickened secondary cell walls [127, 128]. MYB75 is a highly phosphorylation dependent protein with elevated internode expression in maize transcriptome and its role in lianas could be parallel to that of MYB4 [126, 128, 129].

### Downregulated R2R3-MYBs as activators of secondary cell wall synthesis and changers of epidermal properties

Downregulated R2R3-MYBs involved as positive regulators of secondary cell wall synthesis are MYB48, MYB26, MYB39, MYB98, (Table 2, SI Fig. S3). MYB48 appears to play role in xylogenesis but its mechanism of control is unclear containing evidence of alternative splicing through intron-retention [103, 109]. MYB26 positively regulates secondary cell wall biosynthesis in anthers [125]. MYB39 (SUBERMAN) has been shown to be a positive regulator of suberin biosynthesis in Arabidopsis root endodermal tissue [130]. Reduction of suberin in liana leaves maybe a factor in generally low-quality foliage with high turnover as predicted by the LES hypothesis [64, 131].

### R2R3-MYB functions linked to epidermal morphology and plant architecture

MYB98 can be expressed in trichomes and has 83 downstream target genes in synergids [132, 133]. Downregulation of MYB98 maybe a coordinated expression pattern favoring a particular leaf epidermal structure in liana seedlings. This maybe in concert with MYB114 repressor affecting epidermal cell fate and with MYB48 and MYB39 where in grapevine their low expression leads to anthocyanin accumulation in fruit skin [134] (Table 2, SI Figure S3). Branching of the shoots into axillary meristems is controlled by MYB37 and MYB38 also known as regulator of axillary meristems (RAX1 and RAX2) which are positively controlled by a WRKY transcription factor excessive branches 1 (EXB1) [135]. Transcripts of MYB37 and MBY38 are mobile from cell-to-cell [136]. Their downregulation could be a means to control side branching towards unidirectional growth in lianas especially required during the early ontogenetic developmental stages. Shoot meristems can be influenced by a mutant ribosomal protein gene [137]. In our interactome network some of the highest connected nodes were ribosomal proteins such as 40S ribosomal protein S13, 60S ribosomal protein L40, and chloroplast 30S ribosomal protein S5 (Table 2).

## Conclusions

This study explores transcriptomic signatures related to liana leaf properties. Compared to trees, liana leaves generally carry traits colloquially conceptualized as “fast and furious” type life-history strategy associated with quick growth, low cost, high turnover, high capacity for water movement, minimal investment on defense. Our results are in accordance with the LES encapsulating construction costs, rates of carbon fixation and tissue turnover. A set of uniquely expressed enzymes in charge of cell wall building, epicuticular wax synthesis, carbon capture, cell cycle appears to complement with another set of a large number of transcription factors not expressed in lianas. Through comparative transcriptomics of a diverse spectrum of families, we believe we were able to interrogate a wide set of orthologs to contribute towards understanding some of the genetic underpinnings of biology leading to the liana growth form from the leaf perspective. As further sequencing data from additional liana species with more diverse tissue types become available, the genetic basis of this fascinatingly convergent plant form will be more comprehensible.

## Supporting Information

Dryad accession permalink: https://doi.org/10.5061/dryad.n5tb2rbw3 for the following supporting information. **Network file:** A Cytoscape source file (Liana-L5-L2.DRYAD.cys) for the network used in Figure 1. For reproducible results, we recommend selecting Ripple as the visual style and Preset as layout to retain formatting used in the manuscript. **Sequences:** Four fasta formatted files for each species: (a) TRINITY assembled unfiltered transcripts (SPECIES.Trinity.fasta), (b) EnTAP filtered transcripts where non-plant contaminant sequences have been removed (SPECIES.decon.fasta), (c) EnTAP filtered translated/frame-selected full-length protein sequences (SPECIES-fl-protein.fasta), (d) EnTAP filtered full-length transcript nucleotide sequences (SPECIES-fl-nt.fasta).

**SI Dataset S1**

**Spreadsheet**. A spreadsheet (Dataset-Sezen etal Liana.xlsx) containing separate tabs for species list, leaf traits and their PCA scores, QUAST, Bowtie2, BUSCO, gProfiler analysis results, enriched genes, domains and categories.

**SI Figure S1**

**Caption of Fig. S1**. A community representation of transcriptome sequenced trees, shrubs and lianas from Luquillo ForestGEO Forest Dynamics Plot, Puerto Rico. Plant heights reflect approximate mature stature for each species. Lianas are shown with horizontal curves. Non-Luquillo climbers are listed on the upper right.

**SI Figure S2**

**Caption of Fig. S2**. (A) Validation of the de novo assembled transcriptomes via Bowtie2 alignment of trimmed and quality filtered reads. (B) Evaluation for transcriptome completeness using BUSCO analysis. Climbing plant species highlighted in bold. Percentages represent alignment and matching hits.

**SI Figure S3**

**Caption of Fig. S3**. An annotated view of the highlighted nodes within the co-downregulated (light salmon pink boundaries) and co-expressed transcripts (cyan boundaries) with their first shell interactors using the Grapevine interactome (STRING DB v29290). The interactome spans 1107 nodes and 2994 edges. Labels and colors are the same as those in Fig. 1.

**SI Figure S4**

**Caption of Fig. S4**. Ordination of leaf area and specific leaf area traits among lianas and non-lianas in the study. Unit variance scaling was applied to each trait. Principal components were calculated by singular value decomposition with imputation. Prediction ellipses represent 0.95 probability ie. a new observation from the same group expected to fall inside the ellipse. N = 46 data points. Lianas are shown in red. Non-lianas are shown in blue.

## Acknowledgments

We would like to thank Dr. Jill Wegrzyn for being a valuable resource person in bioinformatics and providing access to University of Connecticut’s Xanadu High-Performance Computation Cluster. We would like to thank participants of the 2019 Smithsonian Tropical Research Institute-CTFS ForestGEO annual data analysis workshop in Singapore especially to Dr. Marco Visser for his useful insight and discussion that has helped shape the final version of this manuscript. This work was funded by the US National Science Foundation (DEB-1638488; DEB-1643052).

## Author Contributions

UUS analyzed the data, prepared the figures and wrote the manuscript with input from all authors. UUS NGS SMM conceived and articulated the idea. SJW MNU NGS collected the samples and carried out the molecular lab work. NGS coordinated the sequencing, data curation, analytical verification and provided bioinformatic support. SJW MNU SJD SMM NGS contributed to the conceptualization, writing, review and proofreading, SJD NGS helped with the project coordination, implementation and funding.

