## Supplementary material for "Comparative transcriptomics of tropical woody plants supports fast and furious strategy along the leaf economics spectrum in lianas": SI Supplementary Figures

**This PDF file includes:**

Figs. S1 to S4

**Comparative Transcriptomics of Tropical Woody Plants Supports Fast and Furious Strategy along the Leaf Economics Spectrum in Lianas..**

**U.Uzay Sezen, Samantha J. Worthy, Maria N. Umaña, Stuart J. Davies, Sean McMahon, Nathan G. Swenson.**

**Corresponding Author: Uğur Uzay Sezen**
**.**

**This PDF file includes:**
**Figs. S1 to S4.**

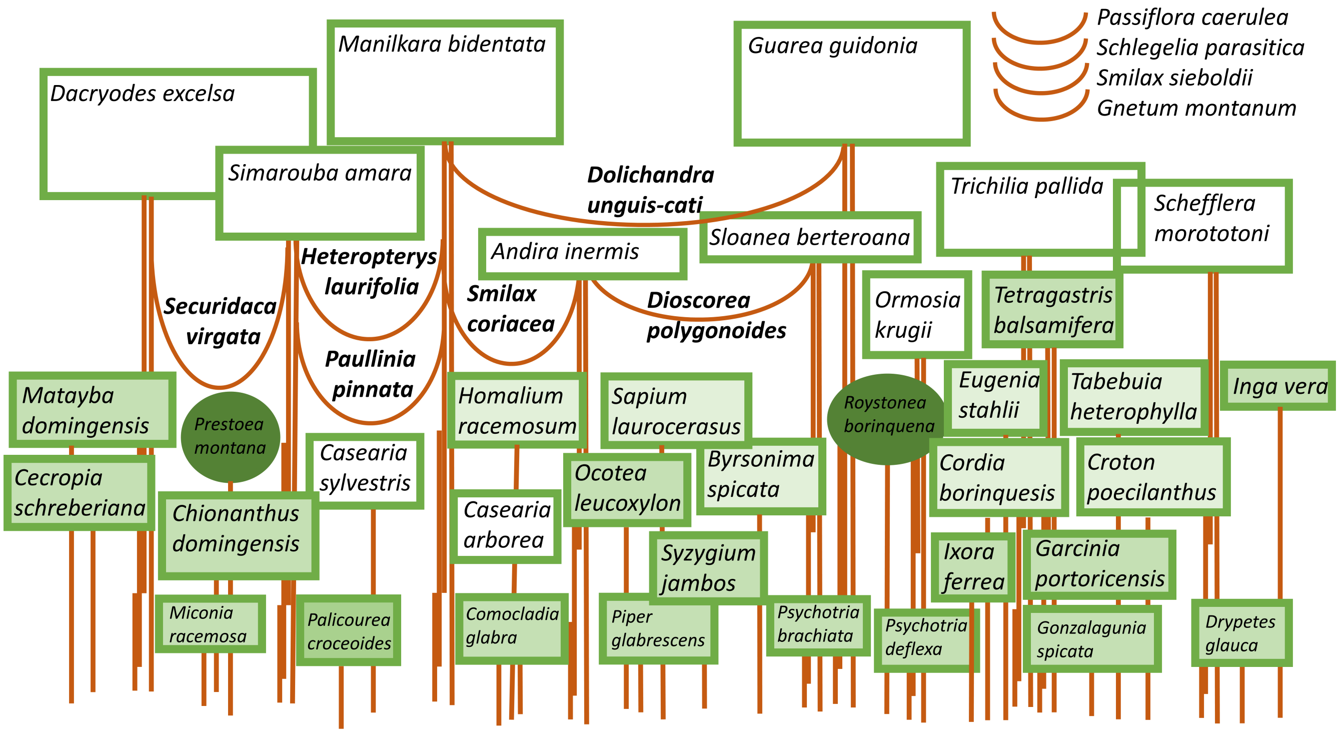


**Fig. S1.** A community representation of transcriptome sequenced trees, shrubs and lianas from Luquillo ForestGEO Forest Dynamics Plot, Puerto Rico. Plant heights reflect approximate mature stature for each species. Lianas are shown with horizontal curves. Non-Luquillo climbers are listed on the upper right. The 16 hectare Puerto Rican ForestGEO plot is located within the Luquillo range which includes the El Yunque National Forest. The Luquillo Experimental Forest (LEF) was established in 1990 and is a Long Term Ecological Research (LTER) site. The biosphere reserve covers 28,000-acres. The plot has been censused 6 times during this publication including 163 species, 165,089 stems and 126,224 stems. Our LEF transcriptome set represented a subset of those found in Luquillo ForestGEO Forest Dynamics Plot (FDP). This cross-sectional representation of LEF species was prepared based on information gathered from the USDA/International Institute of Tropical Forestry 2009 General Technical Report (IITF-GTR-35) by Gary L. Miller and Ariel E. Lugo titled A Guide to the Ecological Systems of Puerto Rico. A full list of species included in our study con be found at the Species tab of Dataset-Sezen_etal_Liana.xlsx deposited in DRYAD accession permalink https://doi.org/10.5061/dryad.n5tb2rbw3

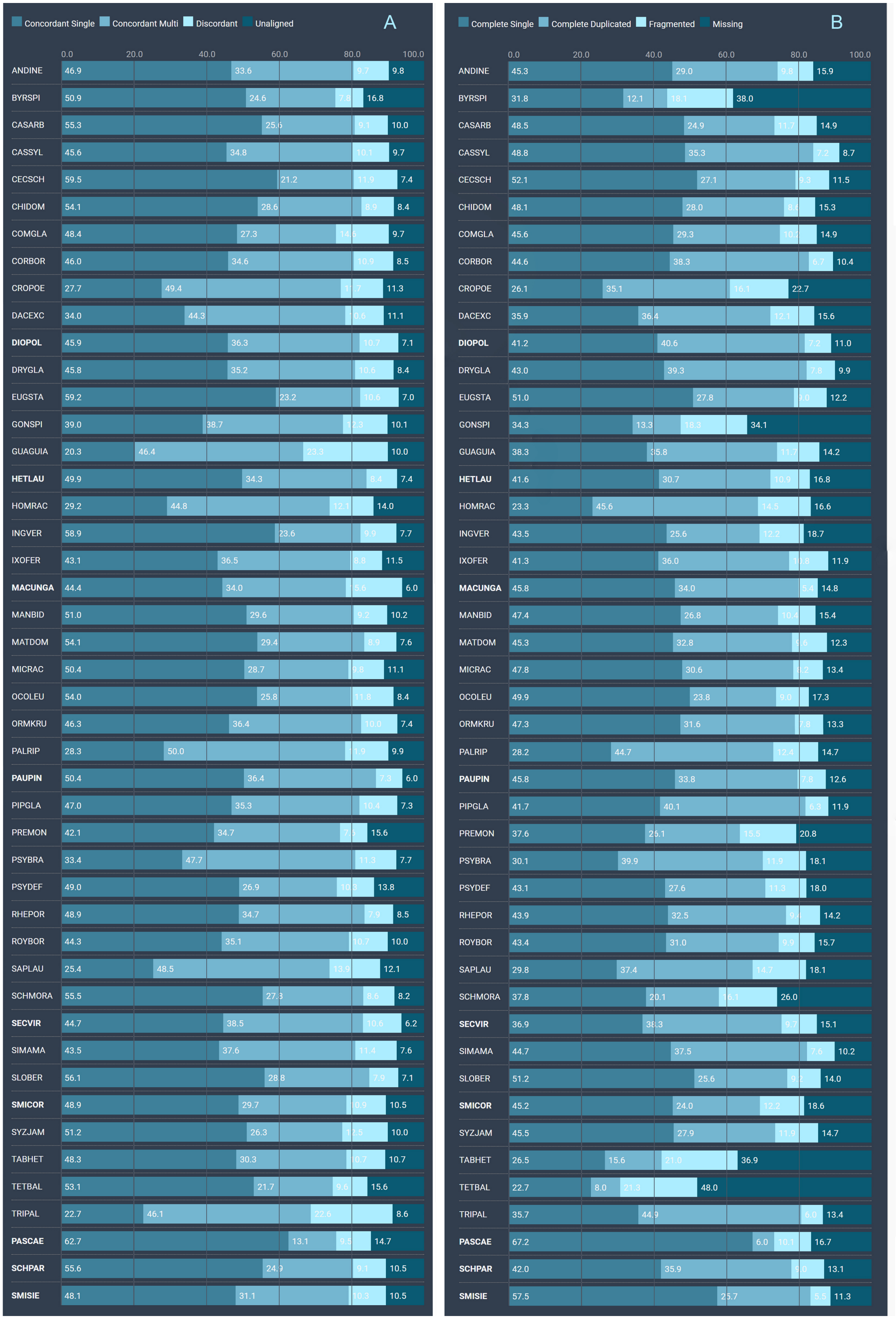


**Fig. S2.** (A) Validation of the de novo assembled transcriptomes via Bowtie2 alignment of trimmed and quality filtered reads. (B) Evaluation for transcriptome completeness using BUSCO analysis. Climbing plant species highlighted in bold. Percentages represent alignment and matching hits.

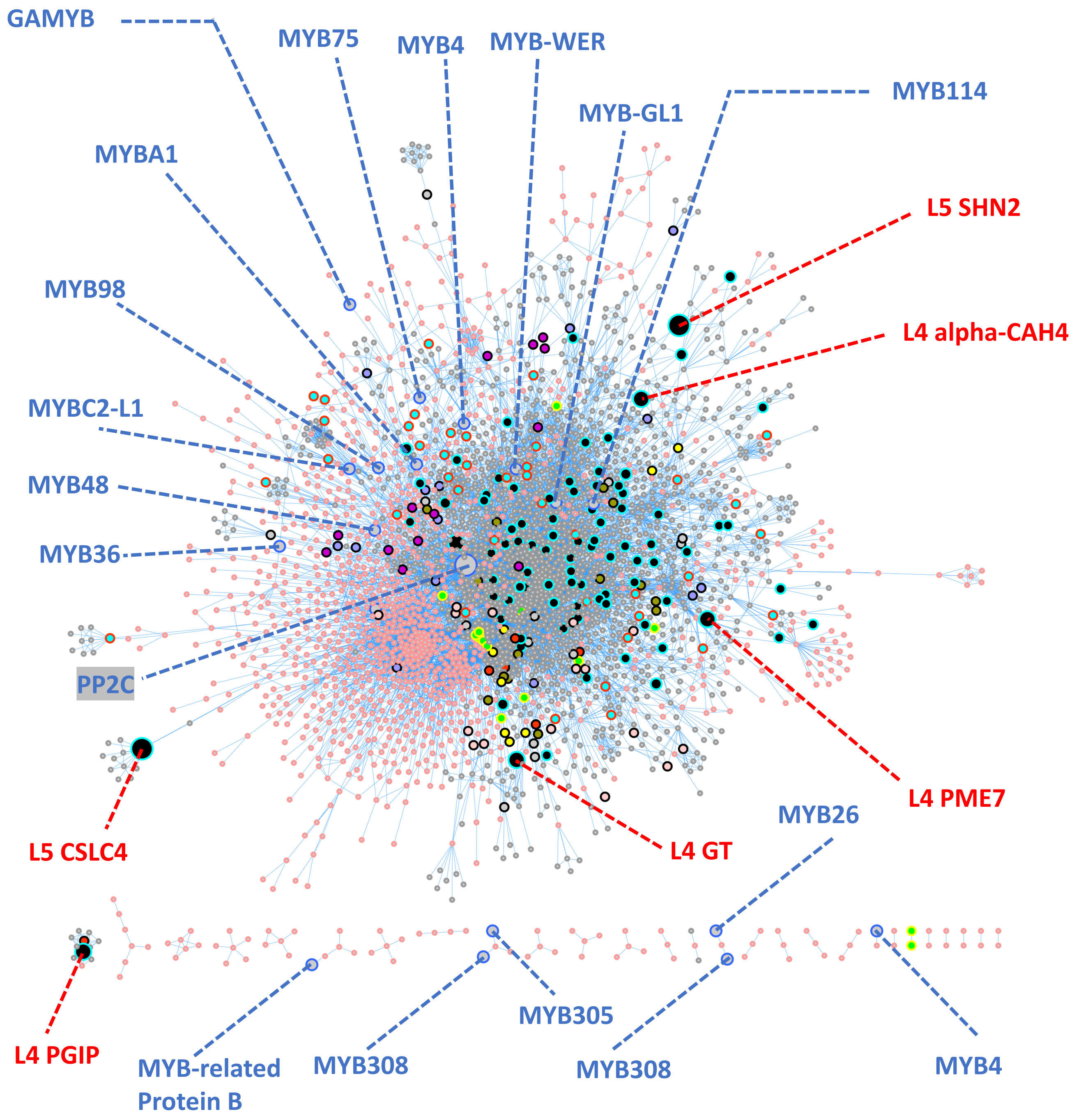


**Fig. S3.** An annotated view of the highlighted nodes within the co-downregulated (light salmon pink boundaries) and co-expressed transcripts (cyan boundaries) with their first shell interactors using the Grapevine interactome (STRING DB v29290). PP2C represents the core of the network with 199 connections. The interactome spans 1107 nodes and 2994 edges. Labels and colors are the same as those in Fig. 1

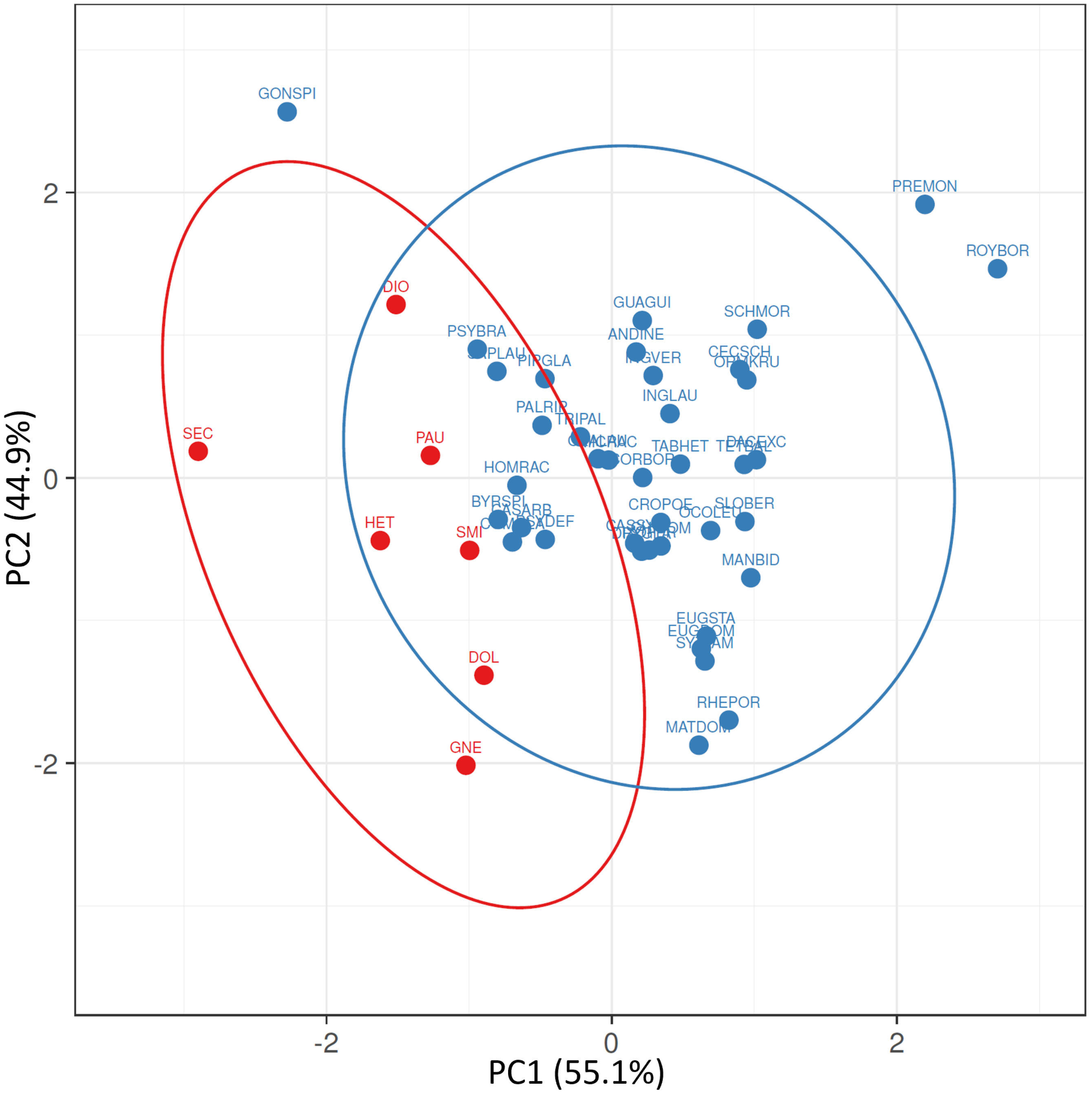


**Fig. S4.** Ordination of leaf area (LA) and specific leaf area (SLA) traits among lianas and non-lianas in the study. Unit variance scaling was applied to each trait. Principal components were calculated by singular value decomposition with imputation. Prediction ellipses represent 0.95 probability ie. a new observation from the same group expected to fall inside the ellipse. N = 46 data points. Lianas are shown in red. Non-lianas are shown in blue. We used Leaf Area (LA) and Specific LA (SLA) trait data from Zambrano et al. 2019 for trees in Luquillo. For *Dolichandra unguis-cati* we used values from Osunkoya et al. (2014). Trait data for *Gnetum montanum* was obtained from the China Trait Database Wang et al. 2018. For the Luquillo vines *Dioscorea polygonoides*, *Heteropterys laurifolia*, *Paullinia pinnata*, *Securidaca virgata*, and *Smilax coriacea* we used unpublished data from the co-authors Samantha J. Worthy and Maria N. Umaña. Ordination of leaf traits was done by the ClustVis webserver. Trait values and calculated PCA scores can be found in a spreadsheet in **SI Dataset S1**. Dryad accession permalink https://doi.org/10.5061/dryad.n5tb2rbw3.
